# Phenotypical mapping of TP53 unique missense mutations spectrum in human cancers

**DOI:** 10.1101/2023.02.06.527255

**Authors:** Lakshay Malhotra, Alankrita Singh, Punit Kaur, Abdul S. Ethayathulla

## Abstract

The p53 tumor suppressor is one of the most mutated genes responsible for tumorigenesis in most human cancers. Out of 29,891 genomic mutations reported in the TP53 Database (https://tp53.isb-cgc.org/), 1297 are identified as unique missense somatic mutants excluding frameshift, intronic, deletion, nonsense, silent, splice and other unknown mutations. we have comprehensively analyzed all these 1297 unique missense mutations and created a phenotypical map based on the distribution of mutants in each domain, the functional state of the protein, and their occurrence in different types of tissues and organs. Our mutation map shows that almost 118 unique missense mutants are reported in the transactivation domain (TADs) and proline-rich domain (PRR), 1,065 in the central DNA-binding domain (DBD), and 113 in the oligomerization (OD) and regulatory domain (RD). Based on the phenotype these 1297 mutations are subdivided into 46 super trans, 491 functional, 315 partially functional, and 415 non-functional mutants. The prevalence of all these mutations was checked in 71 different types of tissues and found the mutant R248Q is reported in 51 types of tissues followed by R175H and R273H in 46 types. The propensity calculation of mutation for each amino acid in p53, showed Proline, Arginine, and Leucine/Glutamic acid are the most frequently mutated residues in the TAD domain, DBD, and TD respectively. We have correlated the impact of these mutations in the structure and function of p53 and highlighted the TP53 unique missense mutants that can be a potential therapeutic drug target with tremendous clinical applications.

## 1. Introduction

According to the Global cancer statistics 2020, the estimated number of cancer cases reported from all over the world is approximately 18,094,716 with breast cancer, lung cancer, prostate, skin, colon, stomach, liver and rectum as the top eight commonly diagnosed cancers [1]. As per reports in the COSMIC database, p53 is the most frequently reported gene in these top 8 cancers like lung (29 %), stomach (37 %), skin (42 %), liver, (29 %) and rectum (60 %) cancers [2]. While in the case of breast cancer the PIK3CA (29 %) gene is at the top followed by the p53 (26 %) gene. Similarly, in colon and prostate cancer, APC (53 %) and LRP1B (37 %) genes respectively are the most frequently mutated genes followed by p53 [3,4]. In normal cells, the transcription factor p53 is negatively regulated by E3 ubiquitin ligase MDM2 to keep it under lower concentration by ubiquitin-mediated proteolysis [5,6]. However, the p53 concentration increases in response to various stresses, such as DNA damage, activation of oncogenes, or hypoxia [7,8]. In response to this genotoxic stress or cellular stress, the TP53 gets activated by posttranslational modifications thereby escaping the MDM2-mediated ubiquitination and translocating inside the nucleus [9]. Inside the nucleus, p53 binds to the appropriate DNA response element and performs its transcription function by activating the genes involved in cell cycle arrest, DNA repair, senescence, or apoptosis [10] depending on cell stress signals. The missense mutation in p53 leads to the elimination of the braking or repair mechanism in a cell leading to tumorigenesis [11]. Mostly the inactivation or deletion of genes coding for crucial proteins like E2F1, RB, WNT signaling regulator APC, BRCA1 is the prime cause of tumorigenesis while in the case of p53, the substitution of a single missense mutation often leads to cancer progression. As per mutations reported in the TP53 database ((https://tp53.isb-cgc.org/), a total of 29,891 mutations are reported so far out of which 2682 are caused by frameshift, 217 intronic, 49 deletion, 2417 non-sense mutations, 21781 missense mutations, 1230 silent mutations, 730 splice mutants, 658 other types of mutations and 127 are unknown [12,13]. Out of the 21,781 missense mutations, 1297 are unique or distinct amino acid residue replacements without including intronic, frameshift, deletion, splicing, multiple mutations, and silent. All the reported 1297 unique somatic mutations are majorly distributed in the three functional domains with 82.11 % (1065) in the DNA binding domain (DBD) followed by 6.01 % in the oligomerization domain of TP53. Studies were reported to understand the impact of these mutations in TP53 and their role in tumor progression and metastasis [11,14,15]. The variants of TP53 either have a loss of function or a new gain of function [16–18]. Various review reports also have described the overall TP53 gene variations, a new gain of functions (GOF), mutation distribution in different populations and clinical applications [3,12,19–22]. In this study, we have performed a detailed analysis of 1297 unique missense somatic mutations of p53 and created a phenotypical map based on the functional state of the mutant, the position of the mutant amino acid in each functional domain and the distribution of mutants in various tissues. Upon analysis only 50 amino acid positions showed no missense mutations out of 393 residues in p53. We also calculated the propensity of mutation of each amino acid residue in p53 by highlighting the important structural and functional regions like DNA contact region, Zinc binding region and post-translational modification (PTM) sites. The analyses give an overall summary of these mutations and their role in cancer progression that can be used for the development of a new drug to restore or inhibit the functions of TP53.

## 2. Methods and datasets

### 2.1. Collection and distribution of p53 unique missense mutations datasets reported in cancers

The unique p53 missense somatic mutants were extracted (excluding frameshift, intronic, deletion, nonsense, silent, splice and other unknown mutations) from *TP53* Database (R20, July 2019): https://tp53.isb-cgc.org) [12,13,19,23]. A dataset was created with these 1,297 unique missense mutations and the dataset was further separated into five datasets based on p53 domains such as Transactivation domain (TAD) (1 to 63), Proline-rich region (PRR) (64 to 91), DNA binding domain (DBD) (92 to 312), Oligomerization or tetramerization domain (OD or TD) (319 to 360) and Regulatory domain (RD) (361-393). Each dataset of different p53 domains was further divided into four phenotypes as described in Kato et al. where the p53 mutants are categorized based on transcription activity using yeast-based functional assay [24]. The mutants with enhanced transcription activity as super trans, with lower activity as partially functional and no activity as non-functional based on the p53 transcription activity on p53 binding elements/promoter such as p21, MDM2, Bax, GADD45, 14-3-3σ, p53AIP1 Noxa and p53R2. Hence, we divided the datasets into super trans, functional, partially functional and non-functional mutants. A heat map was generated based on the phenotypes of p53 using GraphPad Prism 8. Further, the mutants were searched for their prevalence in different human tissues.

### 2.2. Propensity calculation of p53 phenotypical mutants

The propensity of amino acid substitution in each p53 domain was calculated using the following equation

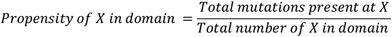

Where X denotes amino acid residue. Using the Expasy Protparam tool, the amino acid composition in each domain was determined and the total number of substitutions present at position X was calculated. Further, the total number of substitutions in each phenotype was calculated using the equation

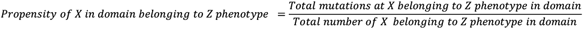

Where Z represents phenotypes (Super trans or functional or partially functional or non-functional). The distribution of all unique missense somatic mutations in 71 different topographies was performed using the collected datasets. The domain-wise topographical distribution of p53 mutations in different phenotypes was performed as described in Kato et al. [24]. The PDBs 3KMD and 1OLG were used to describe the structural features of the p53 DNA binding domain and the tetramerization using Pymol 1.7. The bar diagram and checkers were generated using GraphPad Prism 8.

## 3. Results and Discussion

Of the identified 1,297 unique missense somatic mutations **(Figure 1a, Table 1)** 264 are also found as germline mutations which can pass on to the next generations **(Table 2)**. Upon sub-dividing these distinct missense somatic mutants into each functional domain of TP53, a total of 118 mutants are reported in the transactivation domain (TADs) and proline-rich domain (PRR) (1-91), 1065 in the central DNA-binding domain (DBD) (92-312), and 95 in the oligomerization (OD) and regulatory domain (RD) (319-393), 18 in DBD-OD linker region **(Table 1 and Figure 1b and 1c)**. A total of 46 super trans mutants were found in the whole p53, with 40 in the DBD, 5 in OD, and 1 in TAD **(Table 1) (Figure 1d)**. Surprisingly, no p53 mutant with super trans phenotype was found in the RD and PRR domains. Interestingly, 445 non-functional mutants are reported in p53 out of which 426 are in the DBD, 12 are in OD, 5 in TAD, 1 in PRR and 1 in the DBD-OD linker region **(Table 1, Figure 1d)**. In most cancers, the non-functional and the super-trans mutants promote tumor progression by loss of function and gain of function respectively [18,25]. On the other hand, a total of 491 functional phenotypes (37.86 % of total p53 mutations) are reported in the database, with 315 in the DBD followed by 50, 50, 48, 17 and 11 in OD, TAD, PRR, DBD-OD linker region and RD respectively **(Table 1 and Figure 1d)**. The different phenotypic variants in p53 domains are shown in the p53 mutation heatmap **(Figure 1e)**. While in the case of germline missense mutants, around 238 mutations (90.15 %) reside in the DBD followed by OD (15), TAD (4), CTD (3), DBD-OD linker (2) and PRR (2) domain **(Table 2)**. After dividing the dataset into the functional domain, we analyzed the overall distribution of these mutants in each domain of p53.

**Table 1:** Distribution of p53 missense somatic mutations in different domains.

| **Domains** | **Types of somatic missense mutations** |  |  |  | **Total mutations** | **p53 mutations (%)** |
| --- | --- | --- | --- | --- | --- | --- |
|  | **Super trans** | **Functional** | **Partially functional** | **Non-functional** |  |  |
| **TAD** | 1 | 50 | 9 | 5 | 65 | **5.01** |
| **PRR** | 0 | 48 | 4 | 1 | 53 | **4.08** |
| **DBD** | **40** | **315** | **284** | **426** | **1065** | **82.11** |
| **DBD-OD linker** | 0 | 17 | 0 | 1 | 18 | **1.39** |
| **OD** | 5 | 50 | 11 | 12 | 78 | **6.01** |
| **CTD/ RD** | 0 | 11 | 7 | 0 | 17 | **1.39** |
| **Total** | **46** | **491** | **315** | **445** | **1297** | **-** |

**Table 2:** Distribution of p53 missense germline mutations in different domains.

| **Domains** | **Types of germline missense mutations** |  |  |  | **Total mutations** | **p53 mutations (%)** |
| --- | --- | --- | --- | --- | --- | --- |
|  | **Super trans** | **Functional** | **Partially functional** | **Non-functional** |  |  |
| **TAD** | 0 | 2 | 0 | 2 | 4 | **1.51** |
| **PRR** | 0 | 2 | 0 | 0 | 2 | **0.75** |
| **DBD** | **3** | **28** | **43** | **164** | **238** | **90.15** |
| **DBD-OD linker** | 0 | 2 | 0 | 0 | 2 | **0.75** |
| **OD** | 0 | 8 | 2 | 5 | 15 | **5.68** |
| **CTD/ RD** | 0 | 3 | 0 | 0 | 3 | **1.13** |
| **Total** | **3** | **45** | **45** | **171** | **264** | **-** |

**Figure 1:**
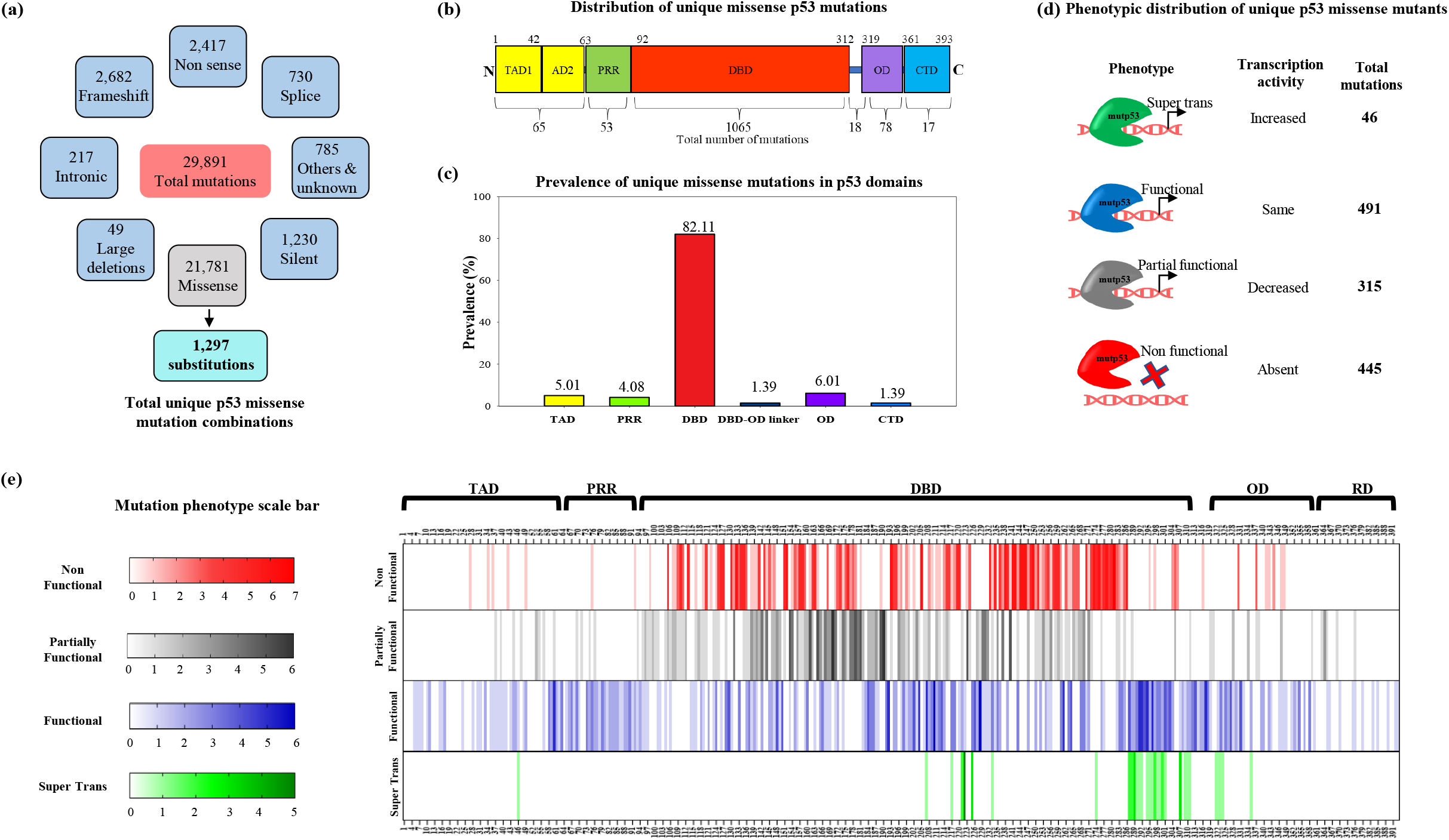
Mutation spectrum of p53. **(a)** Diverse mutants of p53. **(b)** Domain-wise distribution of unique missense mutations. **(c)** Prevalence of unique missense mutations in p53 domains **(d)** Phenotypic distribution of unique p53 missense mutants. **(e)** Heat map of p53 phenotypic mutants.

### 3.1. Transactivation domain (TAD) and Proline-rich region (PRR)

The transactivation domain (TAD) of TP53 can be divided into two distinct domains, TAD1 (1-42) and TAD2 (43 -63) both crucial for tumor suppressor activity of the protein [26,27] **(Figure 2a)**. Each domain has a distinct role or transcriptional requirements related to the p53 tumor suppressor activity [28]. Both TADs is majorly comprised of acidic and/or hydrophobic residues crucial for forming interactions with general transcription factors, Chromatin Modifiers p300 and CBP, and its negative Regulators Mdm2, MdmX, and E1B [29,30]. Similarly, the domain also holds the crucial Posttranslational modifications (PTMs) sites for the activation of p53 **(Table S1)** [9,31–33]. In these domains, a total of 65 unique missense somatic mutations were reported. Out of which 5 are non-functional mutants, 9 are partially functional mutants, 50 are functional mutants and one super trans mutant S46P is reported in this domain **(Table 1) (Figure 2a)**. Out of the five non-functional mutants, P27L, P34L, and P36L indicate that proline at these positions is crucial for the function of TP53 along with D42Y and D49H **(Figure 2a, 2c(iv))**. In the partially functional mutants, the W53C, W53G and F54Y variants are reported as non-functional in combination with other mutants p53^25,26,53,54^ [28] **(Figure 2a and 2c(iii))**. All these hydrophobic residues are important for forming interactions with general transcription factors and regulators [34]. Upon analyzing the propensity of mutation of each amino acid or the phenotypic distribution of the mutations in TAD domains, Proline was found to be the most frequently mutated residue particularly substitution to Leu or Gln **(Figure 2c(ii-iv)**. On the other hand, in the PRR domain a total of 53 mutants were reported, out of which 48 are functional, 4 are partially functional and 1 is non-functional (P75L) **(Table 1 and Figure 2d-f(iii))**. To date, no super transmutation has been reported in the PRR domain. Upon analyzing the propensity of amino acid mutation, Alanine followed by Proline is the most frequently mutated amino residue in the PRR domain with maximum mutations reported as either Pro to Ser or Leu **(Figure 2e and 2f(i))**. In both domains, there are 24 amino acid residue positions with no mutations reported so far **(Figure 2a and 2d)**.

**Figure 2:**
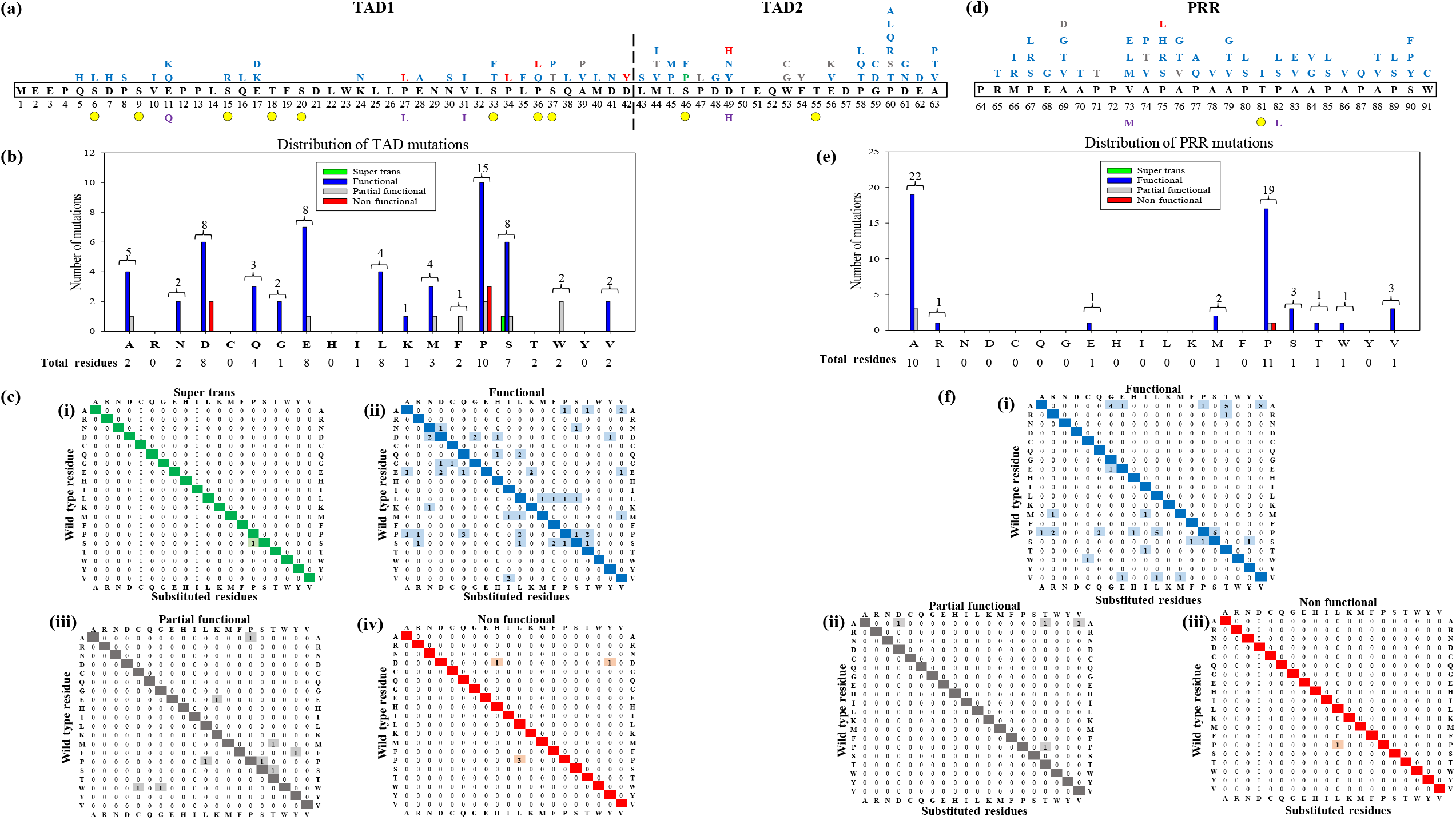
Mapping of missense mutants in transactivation and Proline-rich domains of p53. **(a)** and **(d)** represent the map of somatic mutations in TAD **(a)** and PRR **(d)** domains by different colors based on the phenotype (The residues labeled in red - non-functional phenotype, grey -partial functional phenotype, blue - functional phenotype and green - super trans phenotype). The germline mutant residues are shown in violet color below the wild-type residues. The yellow circle represents PTM sites. **(b)** and **(e)** shows the amino acid residues present in the domain and the propensity of mutations of each residue in TAD and PRR. **(c)** and **(f)** Checkers diagram shows the propensity and frequency of each amino acid substitution in terms of phenotypes (i) super trans, (ii) functional, (iii) partial functional and (iv) non-functional in TAD and PRR.

### 3.2. DNA binding domain or core domain

In the case of the p53 DNA binding domain, 82.11% of the missense mutations are predominantly found in exons 4 to 9 of the p53 gene [12]. Out of 1297, 1065 mutants are reported in DBD with 40 nos’ as super trans, 315 functional, 284 as partially functional phenotypes and 426 as non-functional phenotypes **(Table 1, Figure 3a)**. On the other hand, 238 are found as germline mutations in human cancers **(Table 2)**. Excluding the amino acid Tyr103, all the residues in the DNA binding domain are reported to be mutated in cancer **(Figure 3a)**. The majority of the p53 hotspot mutants are reported in the DNA binding domain due to their presence in the methylated CpG dinucleotides island which is prone to spontaneous mutation by certain environmental carcinogens [35], such as polycyclic aromatic hydrocarbons in tobacco, which preferentially binds to guanines in methylated CpG sites [36,37], and UV irradiation often modifies methylated cytosines [21]. To assess the propensity of mutation in each amino acid residue in the DNA binding domain we divided the residues based on their location in the domain such as the DNA contact region, Zinc binding region and conformational mutants that lead to the loss of thermostability of p53 **(Figure 4a-c)**.

**Figure 3:**
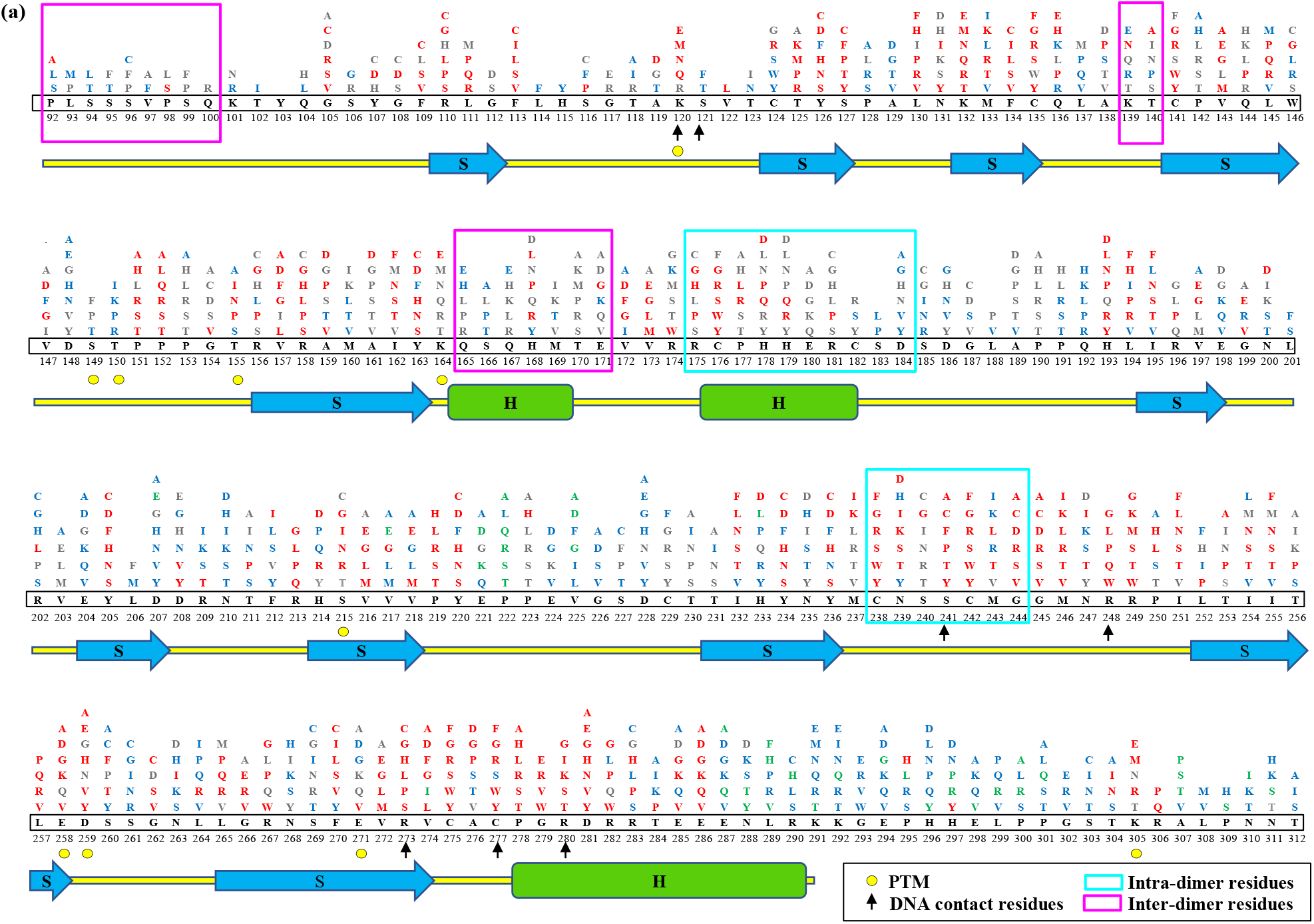
Mapping of missense mutants in the p53 DNA binding domain. The different phenotypic mutants are shown along with the secondary structure of the p53 DNA binding domain. The mutant residues are highlighted on the top of wt residues with different colors based on the phenotype of the mutations. (The residues labeled in red - non-functional phenotype, grey -partial functional phenotype, blue - functional phenotype and green - super trans phenotype). S-beta strand (Blue arrow), H-alpha helix (green cylinder) and the unstructured region (loops and coils) in a yellow line.

**Figure 4:**
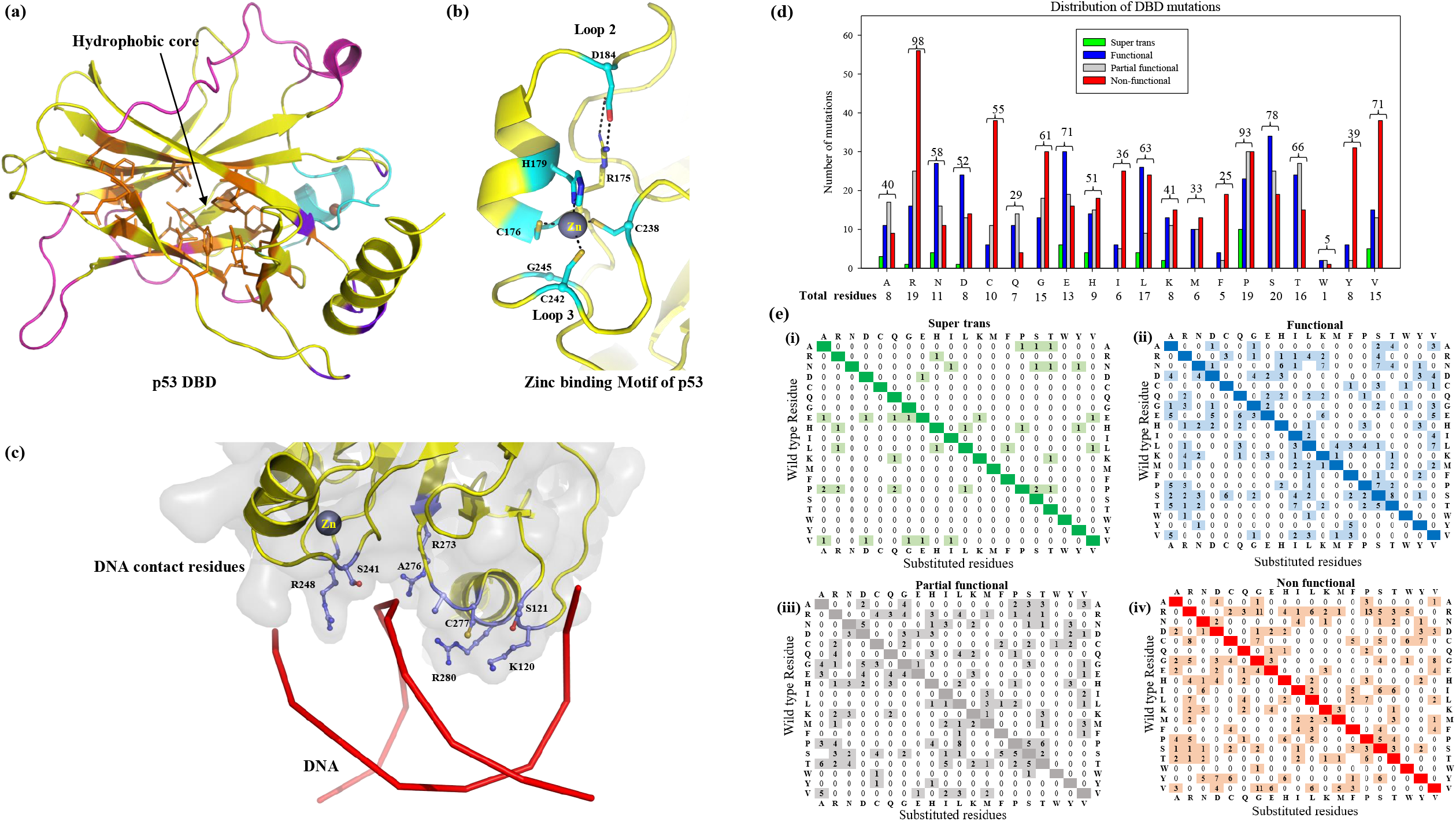
Structure of p53 DNA binding domain and the mutant distribution. **(a)** p53 DBD cartoon highlighting Zinc binding motif in cyan, DNA contact region in blue, hydrophobic core residues are shown in orange and dimerization region in magenta. **(b)** Represents the Zinc binding motif and the residues involved in Zinc coordination are shown in cyan. **(c)** The DNA contact residues of p53DBD are shown in blue. **(d)** Phenotypical distribution of mutations for each type of amino acid in DBD. **(e)** Phenotype-wise checkers diagrams (**(i)** super trans functional, **(ii)**, partial functional **(iii)** and non-functional **(iv)**) represent the propensity of a particular amino acid to get mutated in p53 DBD.

TP53 recognizes DNA with a loop-sheet-helix motif and the residues involved in direct interactions with DNA are Lys120, Ser121, Ser241, Arg248, Arg273, Ala276, Cys277 and Arg280 **(Figure 4c)**. Interestingly all the arginine residues which are involved in DNA contact were found to be mutated and all the mutants belong to the Non-functional phenotype **(Figure 3a, 4e(iv))**. On the other hand, the mutations reported at the Lys120 position are majorly non-functional except for K120R which turns out to be partially functional **(Figure 3a)**. Surprisingly, the mutations occurring at Ser121 (S121F, S121T) and Ala276 (A276S and A276T) belong to the functional category **(Figure 3a)**. The DNA contact mutants are considered as first hotspot mutants. Remarkably, apart from DNA contact residues, substituting any residue that is part of the loop-sheet-helix motif makes TP53 a non-functional protein. The TP53 recognizes the promoter sequence as a tetramer (dimer of dimers) and the dimerization interface in the DNA binding domain is the 3^rd^ hot spot mutation site **(Figure 3a and 4a)**. Dimerization is crucial for DNA binding and regulating the transcription of genes. The dimer interface consists of a Zinc binding motif formed by two loops (L2 and L3) where the zinc atom forms a tetrahedral coordination bond with C176 and H179 of the L2 loop and C238 and C242 of the L3 loop **(Figure 4b)**. TP53 is extremely unstable in absence of zinc and it is predicted to be unfolded state in the cell [38]. Out of these four amino acid residues, C176 and H179 are the 7^th^ and 8^th^ most frequently mutated residues in p53 **(Table 3)**. Apart from these zinc-coordination residues, the p53R175 and G245 are also reported as high-frequency hotspot mutants in almost all cancers **(Table 3)**. The R175 and G245 are located near the zinc-binding site in the DBD dimer and DNA binding interface maintaining the stability of the zinc-binding motif **(Figure 4b)**. Arg175 forms a salt-bridge interaction with Asp184 and G245 is part of the 3_10_ helix which holds the Zinc ion **(Figure 4b)**. Mutations of Arg and Gly lead to non-functional proteins **(Figures 3a)**. Apart from DNA-contact and Zinc binding mutants, several structural mutants are also reported to cause non-functional phenotypes. The mutations lead to either distortion in the DNA binding surface or to cause formation of water-accessible crevices or lead variations in dimer-dimer interacting interfaces involved in tetramerization **(Figure 3a and 4a)**. Many structural mutants are considered therapeutic targets rescued by small molecule drugs to restore protein function in tumors, especially Y220C [39–44] **(Table 3)**. The other structural mutants are the residues in the hydrophobic core of the DBD **(Figure 4a)**. The hydrophobic core holds and stabilized the domain upon DNA binding. The core is formed by the residues F109, L111, F113, C124, Y126, M133, C135, C141, V143, L145, V157, A159, I195, V197, V216, V218, I232, Y234, Y236, T253, I255, L257, F270 and V274 **(Figure 4a)**. The mutations in these residues lead to mostly non-functional proteins **(Figure 3a)**. Interestingly, upon analyzing the phenotypic distribution of all amino acid residues in the DNA binding domain, a total of 98 mutations are reported for Arginine (19 in the domain) **(Figures 4d and 4e)**. The 19 residues are found to be most frequently reported with an average of 5.2 mutations per arginine in DBD and the maximum number of mutants are non-functional **(Figure 4d and 4e(iv))**. Out of all arginine mutants, only Arg → His (1) mutation leads to super trans phenotype **(Figure 3a and 4e(i))**. On the other hand, Arg → Cys (3), Arg → Gly or His or Ile (1), Arg → Leu or Ser (4), and Arg → Lys (2) **(Figure 4e(ii))** are functional. Whereas in the case of p53 mutants elucidating partially functional phenotype, Arg → Cys or Gly or Leu or Ser (4), Arg → Gln or His (3), and Arg → Met or Pro or Thr (1) **(Figure 4e(iii))**. The non-functional mutants are Arg → Pro (13), Arg → Gly (11), Arg → Leu (6), Arg → Ser or Trp 5 times, Arg → His 4 times, Arg → Thr or Gln 3 times each, Arg → Ile or Met (1). Similarly, the analysis was performed for other amino acid residues shown in **Figure 4e**. It is also reported that post-translational modifications in the DBD enhance the DNA binding affinity of p53 **(Table S1)**. The domain was reported to have 10 potential PTM sites [9,45] and mutations are observed at these sites **(Figure 3a)**.

**Table 3:** Top 10 most frequently mutated residues of p53.

| S.No. | Amino Acid Residue | Somatic | Germline | Type |
| --- | --- | --- | --- | --- |
| 1 | R248 | G, L, P, Q, W | L, Q, W | DNA contact |
| 2 | R273 | C, G, H, L, P, S | C, G, H, L, S | DNA contact |
| 3 | R175 | C, G, H, L, P, S | C, G, H, L | Zinc binding motif |
| 4 | G245 | A, C, D, R, S, V | C, D, S, V | Zinc binding motif |
| 5 | R249 | G, K, M, S, T, W | K, T | DNA contact |
| 6 | Y220 | C, D, F, H, N, S | C, S | Structural |
| 7 | C176 | F, G, R, S, W, Y | G, Y | Zinc binding |
| 8 | H179 | D, L, N, P, Q, R, Y | D, Q, R, Y | Zinc binding |
| 9 | V157 | A, D, F, G, I, L | - | Structural |
| 10 | M237 | I, K, L, R, T, V | - | Structural |

### 3.3. Mutants in the p53 tetramerization (TD) and the regulatory domain

The C-terminus of TP53 consists of tetramerization (TD) and the regulatory domain (RD). The oligomeric form of p53 is crucial for its transcription function and it is reported that the DNA binding affinity increases several folds upon tetramerization [46]. The post-translational modifications (23 PTM sites) in the C-terminal domain also dictate many functional roles of p53 in protein-protein interactions [47] **(Figure 5a and 6a and Table S1)**. The TD also has important sequence motifs like nuclear localization signal (NLS1) and nuclear export signal (NES) **(Figure 5a)**.

**Figure 5:**
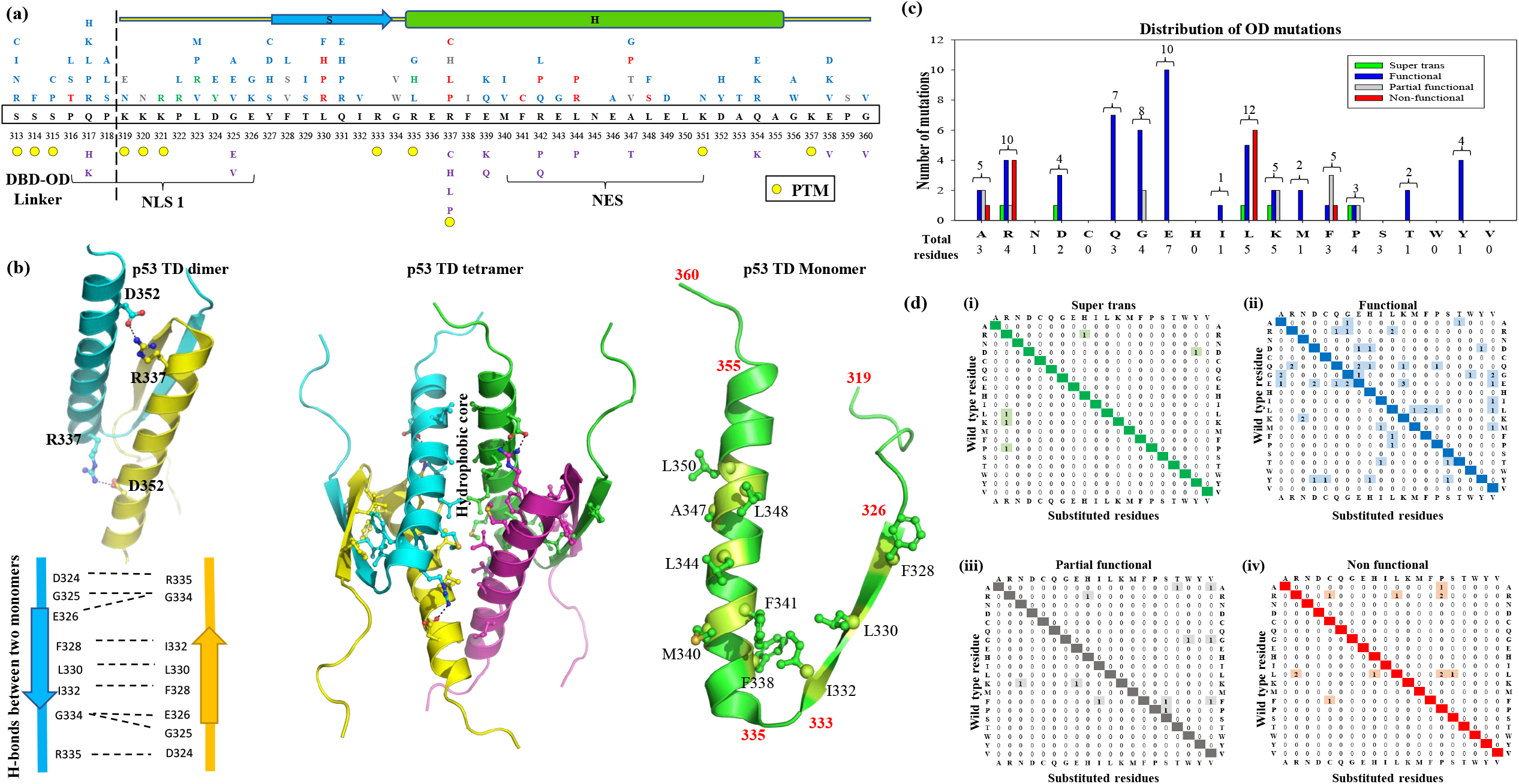
Mutation mapping and distribution of p53 oligomerization mutations. **(a)** The mutation map represents somatic mutations in the Oligomerization or tetramerization domain by different colors based on the phenotype (The amino acid single letter code with red color represents non-functional phenotype, grey color represents partial functional phenotype, blue color denotes functional phenotype and green color represents super trans phenotype). The germline mutations are also shown in violet color below the wild-type residues. **(b)** Cartoon diagram OD of p53 showing the structure of p53 tetramer formed by OD (the monomers are shown in cyan, yellow, neon, and magenta), the residues involved in the formation of the dimer (cyan and yellow), and the hydrophobic residues involved in the formation of a dimer of dimer or tetramerization of p53 (green). **(c)** Represent the propensity of mutations of amino acid residues in p53 TD. **(d)** Phenotype-wise checkers diagrams (**(i)** functional and **(ii)** partial functional) representing the propensity of a particular amino acid to get mutated in p53 DBD, where the Y-axis denotes the wild-type residue whereas the X-axis denotes the total number of different combinations present.

The active TP53 forms a tetramer, a dimer of dimers where the TD plays an important role **(Figure 5b)** [48,49]. Structurally the monomeric tetramerization domain (319-360) of p53 consists of a β-strand (residues 326 to 333) and an α-helix (residues 335 to 355) connected by a flexible residue G334 [50] **(Figure 5a, 5b)**. Two monomers interact with each other through an antiparallel double-stranded β-sheet and antiparallel helices to form a double-helical bundle **(Figure 5b)**. Four monomers of p53TD form a V-shaped fold with a hydrophobic core formed by F328, L330, I332, F338, M340, F341, L344, A347, L348 and L350 **(Figure 5b)** from each monomer. The four monomers are also held by four salt bridges formed by R337 and D352 from different monomers **(Figure 5b)**. The mutation at positions F341 and L344 are non-functional while L330, R337, R342, A347 and L348 are reported to be mostly non-functional highlighting the importance of these hydrophobic residues in stabilizing the domain and p53 **(Figure 5a)**. The OD domain was reported to have a second maximum number of mutants in p53, accounting for 78 unique missense mutants in human cancers **(Table 1)**. Among these mutants, 50 are functional, 12 non-functional mutants, 11 are partially functional and 5 are super trans **(Figure 5a, Table 1)**. Upon analyzing the propensity of amino acid mutations in this domain, Leucine followed by Glutamic acid is the most frequently mutated residue in the TD domain (**Figures 5c and 5d)**. The mutations at the DBD-OD linker region between 313 to 318 were mostly reported as functional and this region contains 3 PTM sites **(Figure 5a)**.

Finally, the regulatory domain (RD) is known to be the least mutated domain in p53 to date with 18 mutations, out of which 11 are functional and 7 are partially functional. RD is the only domain that lacks non-functional and super trans mutation **(Table 1, Figure 6a-c)**. The most important feature is the 12 post-translational modification sites particularly phosphorylation and acetylation at K370, K381 and K382 which converts the p53 into active upon cellular stress **(Figure 6a, Table S1)**. The RD is also proposed to regulate the DNA binding of p53 and K373, K381 and K382 are predicted to be involved in DNA binding [51]. The RD also has NLS2 and NLS3 regions where the mutations were reported at these sites but found to have functional phenotype **(Figure 6a)**. Upon analyzing the propensity of amino acid mutations in this domain, Serine and Alanine are the most frequently mutated residues in the RD domain (**Figures 6b and 6c)**.

**Figure 6:**
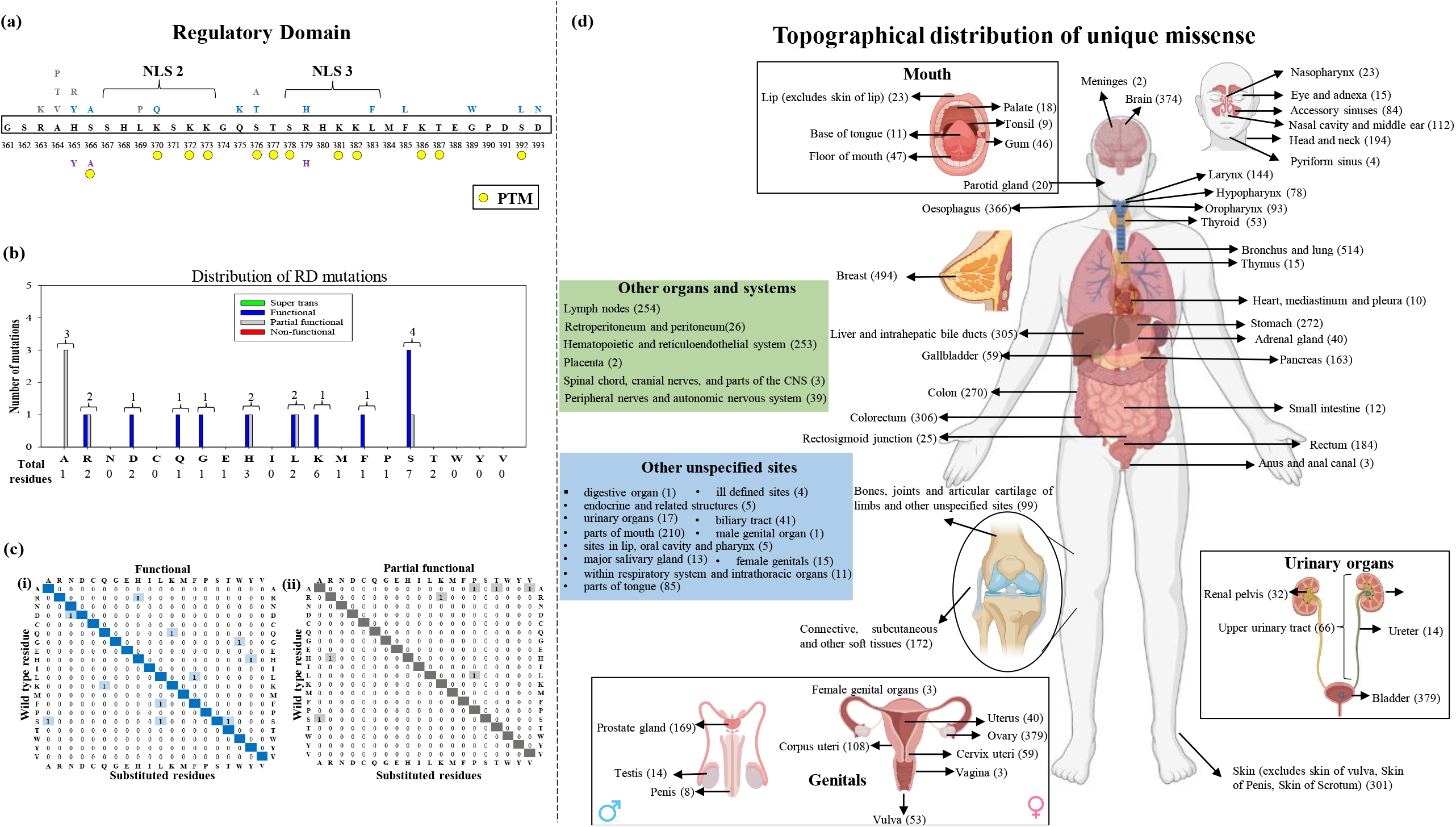
Mutation mapping and distribution of p53 Regulatory domain: **(a)** The mutation map represents somatic mutations in the regulatory domain by different colors based on the phenotype (The amino acid single letter code grey color represents partial functional phenotype; blue color denotes functional phenotype). The germline mutations are also shown in violet color below the wild-type residues. **(b)** Represent the propensity of mutations of amino acid residues in p53 RD. **(c)** Phenotypical distribution of mutations at each type of amino acid of p53 RD shown by checkers diagrams **(i)** functional and (**ii)** partially functional mutants. **(d)** Represents topographical distribution of p53 unique missense mutants in 71 different tissue topographies. The total number of mutants observed in each tissue type is given in brackets.

### 3.4. Distribution of TP53 mutants in the tissues

The phenotypic distribution of all the mutants from different domains in various tissues showed that the variants are reported in almost 71 different types of tissues **(Figure 6d) (Table S2)**. Out of these, the DBD mutants are reported in 71 different types of tissues while DBD-OD-linker region mutants are in 18 types, OD mutants in 29 types, RD in 13 types and TAD-PRR domain mutants in 33 types of tissues **(Table S3-7)**. Out of these R248Q was reported in 51 different types of tissues, R175H and R273H in 46 types, and mutants R248W, R273C, R282W, G245S, Y220C, E285K and C176F were reported in more than 30 types of tissues **(Table 4)**. The phenotypical distribution of these mutants is shown in **Supplementary Table S3-7**.

**Table 4:** List of most dominant missense somatic mutants observed in ≥ 25 different types of human cancers.

| **Mutant** | **Total number of tissues where the mutant is reported** |
| --- | --- |
| R248Q | 51 |
| R175H | 46 |
| R273H | 46 |
| R248W | 43 |
| R273C | 41 |
| R282W | 40 |
| G245S | 36 |
| Y220C | 33 |
| E285K | 33 |
| C176F | 32 |
| H179Y | 31 |
| Y205C | 30 |
| G245D | 30 |
| R249S | 28 |
| V173M | 27 |
| M237I | 26 |
| G266E | 26 |
| R280T | 26 |
| P151S | 25 |
| Y163C | 25 |
| Y234C | 25 |
| G244D | 25 |
| E258K | 25 |
| V272M | 25 |

## 4. Conclusion

In several cancers, the heterozygous state where both wildtype and mutants co-exist, often found that the mutant protein is in the dominant state [52–54]. Our map shows the list of all potential p53 mutants which have been found in numerous human cancers till date. In addition, the map also describes the list of all super trans, functional, partial functional and non-functional p53 mutations. Our analysis gives an overall idea of all potential therapeutic targets in p53 that can be used to identify drugs, small molecule chaperones or small peptides to restore its wild-type function or to eliminate mutant p53 by immunotherapies to prevent the progression of tumor cells.

## Abbreviations

DBD: DNA binding domain;
NES: Nuclear export signal;
NLS: Nuclear localization signal;
OD: Oligomerization domain;
PDB: Protein data bank;
PTM: Post-translational modification;
PRR: Proline-rich region;
RD: Regulatory domain;
TAD: Transactivation domain;
wt: wild type

## CRediT authorship contribution statement

**L.M:** Conceptualization, Methodology, Validation, Formal analysis, Investigation, Data Curation, Graphics, Writing, Review & Editing, and Visualization. **A.S:** Validation, Formal analysis, Data Curation. **P.K:** Data Curation. **A.S.E:** Methodology, Validation, Formal analysis, Investigation, Data Curation, Writing - Original Draft, Writing - Review & Editing, and Visualization.

## Ethical approval

The article does not involve any study with human participants or animals to be performed by any of the authors.

## Acknowledgments

**LM**, and **AS** are thankful to the Department of Biotechnology (DBT) for providing the Junior Research Fellowship. **ASE** is thankful to the Indian Council of Medical Research (ICMR) for the financial assistance of the project (grant ID No: 5/13/72/2020/NCD-III) and the All India Institute of Medical Sciences (AIIMS) for providing the facility.

## Data Availability Statements

The data underlying this article will be shared on reasonable request to the corresponding author.

## References

1. Sung H, Ferlay J, Siegel RL, et al. Global Cancer Statistics 2020: GLOBOCAN Estimates of Incidence and Mortality Worldwide for 36 Cancers in 185 Countries. CA Cancer J Clin 2021; 71:209–249

2. Tate JG, Bamford S, Jubb HC, et al. COSMIC: the Catalogue Of Somatic Mutations In Cancer. Nucleic Acids Research 2019; 47:D941–D947

3. Zhu G, Pan C, Bei J-X, et al. Mutant p53 in Cancer Progression and Targeted Therapies. Frontiers in Oncology 2020; 10:

4. Kandoth C, McLellan MD, Vandin F, et al. Mutational landscape and significance across 12 major cancer types. Nature 2013; 502:333–339

5. Hock AK, Vousden KH. The role of ubiquitin modification in the regulation of p53. Biochimica et Biophysica Acta (BBA) - Molecular Cell Research 2014; 1843:137–149

6. Giaccia AJ, Kastan MB. The complexity of p53 modulation: emerging patterns from divergent signals. Genes Dev. 1998; 12:2973–2983

7. Lavin MF, Gueven N. The complexity of p53 stabilization and activation. Cell Death Differ 2006; 13:941–950

8. Ljungman M. Dial 9-1-1 for p53: Mechanisms of p53 Activation by Cellular Stress. Neoplasia 2000; 2:208–225

9. Meek DW, Anderson CW. Posttranslational Modification of p53: Cooperative Integrators of Function. Cold Spring Harb Perspect Biol 2009; 1:a000950

10. Zilfou JT, Lowe SW. Tumor Suppressive Functions of p53. Cold Spring Harb Perspect Biol 2009; 1:a001883

11. Malhotra L, Singh A, Kaur P, et al. Comprehensive omics studies of p53 mutants in human cancer. Briefings in Functional Genomics 2022; elac015

12. Bouaoun L, Sonkin D, Ardin M, et al. TP53 Variations in Human Cancers: New Lessons from the IARC TP53 Database and Genomics Data. Human Mutation 2016; 37:865–876

13. M O, R E, M H, et al. The IARC TP53 database: new online mutation analysis and recommendations to users. Human mutation 2002; 19:

14. Tang Q, Su Z, Gu W, et al. Mutant p53 on the Path to Metastasis. Trends in Cancer 2020; 6:62–73

15. Powell E, Piwnica-Worms D, Piwnica-Worms H. Contribution of p53 to metastasis. Cancer Discov 2014; 4:405–414

16. Oren M, Rotter V. Mutant p53 Gain-of-Function in Cancer. Cold Spring Harbor Perspectives in Biology 2010; 2:

17. Zhang C, Liu J, Xu D, et al. Gain-of-function mutant p53 in cancer progression and therapy. Journal of Molecular Cell Biology 2020; 12:674–687

18. Alvarado-Ortiz E, de la Cruz-López KG, Becerril-Rico J, et al. Mutant p53 Gain-of-Function: Role in Cancer Development, Progression, and Therapeutic Approaches. Frontiers in Cell and Developmental Biology 2021; 8:

19. de Andrade KC, Lee EE, Tookmanian EM, et al. The TP53 Database: transition from the International Agency for Research on Cancer to the US National Cancer Institute. Cell Death Differ 2022; 29:1071–1073

20. Olivier M, Hollstein M, Hainaut P. TP53 Mutations in Human Cancers: Origins, Consequences, and Clinical Use. Cold Spring Harb Perspect Biol 2010; 2:

21. Parkin DM, Mesher D, Sasieni P. 13. Cancers attributable to solar (ultraviolet) radiation exposure in the UK in 2010. Br J Cancer 2011; 105 Suppl 2:S66–69

22. Sabapathy K, Lane DP. Therapeutic targeting of p53: all mutants are equal, but some mutants are more equal than others. Nat Rev Clin Oncol 2018; 15:13–30

23. Leroy B, Fournier JL, Ishioka C, et al. The TP53 website: an integrative resource centre for the TP53 mutation database and TP53 mutant analysis. Nucleic Acids Res 2013; 41:D962–969

24. Kato S, Han S-Y, Liu W, et al. Understanding the function–structure and function–mutation relationships of p53 tumor suppressor protein by high-resolution missense mutation analysis. PNAS 2003; 100:8424–8429

25. Rivlin N, Brosh R, Oren M, et al. Mutations in the p53 Tumor Suppressor Gene. Genes Cancer 2011; 2:466–474

26. Baral A, Asokan A, Bauer V, et al. Chemical synthesis of transactivation domain (TAD) of tumor suppressor protein p53 by native chemical ligation of three peptide segments. Tetrahedron 2019; 75:703–708

27. Lee CW, Martinez-Yamout MA, Dyson HJ, et al. Structure of the p53 transactivation domain in complex with the nuclear coactivator binding domain of CBP. Biochemistry 2010; 49:9964–9971

28. Brady CA, Jiang D, Mello SS, et al. Distinct p53 Transcriptional Programs Dictate Acute DNA Damage Responses and Tumor Suppression. Cell 2011; 145:571–583

29. Collavin L, Lunardi A, Del Sal G. p53-family proteins and their regulators: hubs and spokes in tumor suppression. Cell Death Differ 2010; 17:901–911

30. Fernandez-Fernandez MR, Sot B. The relevance of protein–protein interactions for p53 function: the CPE contribution. Protein Eng Des Sel 2011; 24:41–51

31. Gu B, Zhu W-G. Surf the Post-translational Modification Network of p53 Regulation. Int J Biol Sci 2012; 8:672–684

32. Che Z, Sun H, Yao W, et al. Role of post-translational modifications in regulation of tumor suppressor p53 function. Frontiers of Oral and Maxillofacial Medicine 2020; 2:

33. Dai C, Gu W. p53 post-translational modification: deregulated in tumorigenesis. Trends in Molecular Medicine 2010; 16:528–536

34. Raj N, Attardi LD. The transactivation domains of the p53 protein. Cold Spring Harbor Perspectives in Medicine 2017;

35. Rosendahl Huber A, Van Hoeck A, Van Boxtel R. The Mutagenic Impact of Environmental Exposures in Human Cells and Cancer: Imprints Through Time. Front Genet 2021; 12:760039

36. Peto R, Darby S, Deo H, et al. Smoking, smoking cessation, and lung cancer in the UK since 1950: combination of national statistics with two case-control studies. BMJ 2000; 321:323–329

37. Pfeifer GP, Denissenko MF, Olivier M, et al. Tobacco smoke carcinogens, DNA damage and p53 mutations in smoking-associated cancers. Oncogene 2002; 21:7435–7451

38. Blanden AR, Yu X, Blayney AJ, et al. Zinc shapes the folding landscape of p53 and establishes a pathway for reactivating structurally diverse cancer mutants. eLife 2020; 9:e61487

39. Malhotra L, Sharma S, Hariprasad G, et al. Mechanism of apoptosis activation by Curcumin rescued mutant p53Y220C in human pancreatic cancer. Biochim Biophys Acta Mol Cell Res 2022; 1869:119343

40. Malhotra L, Goyal HKV, Jhuria S, et al. Curcumin rescue p53Y220C in BxPC-3 pancreatic adenocarcinomas cell line: Evidence-based on computational, biophysical, and in vivo studies. Biochimica et Biophysica Acta (BBA) - General Subjects 2021; 1865:129807

41. Raghavan V, Agrahari M, Gowda DK. Virtual screening of p53 mutants reveals Y220S as an additional rescue drug target for PhiKan083 with higher binding characteristics. Computational Biology and Chemistry 2019; 80:398–408

42. Joerger AC, Ang HC, Fersht AR. Structural basis for understanding oncogenic p53 mutations and designing rescue drugs. Proceedings of the National Academy of Sciences 2006; 103:15056–15061

43. Bauer MR, Krämer A, Settanni G, et al. Targeting Cavity-Creating p53 Cancer Mutations with Small-Molecule Stabilizers: the Y220X Paradigm. ACS Chem. Biol. 2020; 15:657–668

44. Bauer MR, Jones RN, Tareque RK, et al. A structure-guided molecular chaperone approach for restoring the transcriptional activity of the p53 cancer mutant Y220C. Future Medicinal Chemistry 2019; 11:2491–2504

45. Gu B, Zhu W-G. Surf the Post-translational Modification Network of p53 Regulation. Int J Biol Sci 2012; 8:672–684

46. Lubin DJ, Butler JS, Loh SN. Folding of tetrameric p53: oligomerization and tumorigenic mutations induce misfolding and loss of function. J. Mol. Biol. 2010; 395:705–716

47. Laptenko O, Shiff I, Freed-Pastor W, et al. The p53 C Terminus Controls Site-Specific DNA Binding and Promotes Structural Changes within the Central DNA Binding Domain. Molecular Cell 2015; 57:1034–1046

48. Clore GM, Omichinski JG, Sakaguchi K, et al. High-Resolution Structure of the Oligomerization Domain of p53 by Multidimensional NMR. Science 1994; 265:386–391

49. Chène P. The role of tetramerization in p53 function. Oncogene 2001; 20:2611–2617

50. Jeffrey PD, Gorina S, Pavletich NP. Crystal structure of the tetramerization domain of the p53 tumor suppressor at 1.7 angstroms. Science 1995; 267:1498–1502

51. Weinberg RL, Freund SMV, Veprintsev DB, et al. Regulation of DNA Binding of p53 by its C-terminal Domain. Journal of Molecular Biology 2004; 342:801–811

52. Milner J, Medcalf EA. Cotranslation of activated mutant p53 with wild type drives the wild-type p53 protein into the mutant conformation. Cell 1991; 65:765–774

53. Milner J, Medcalf EA, Cook AC. Tumor suppressor p53: analysis of wild-type and mutant p53 complexes. Mol Cell Biol 1991; 11:12–19

54. Cuddihy AR, Jalali F, Coackley C, et al. WTp53 induction does not override MTp53 chemoresistance and radioresistance due to gain-of-function in lung cancer cells. Mol Cancer Ther 2008; 7:980–992

