## Supplementary material for "Phenotypical mapping of TP53 unique missense mutations spectrum in human cancers": Figure S1

### Germline DBD mutants

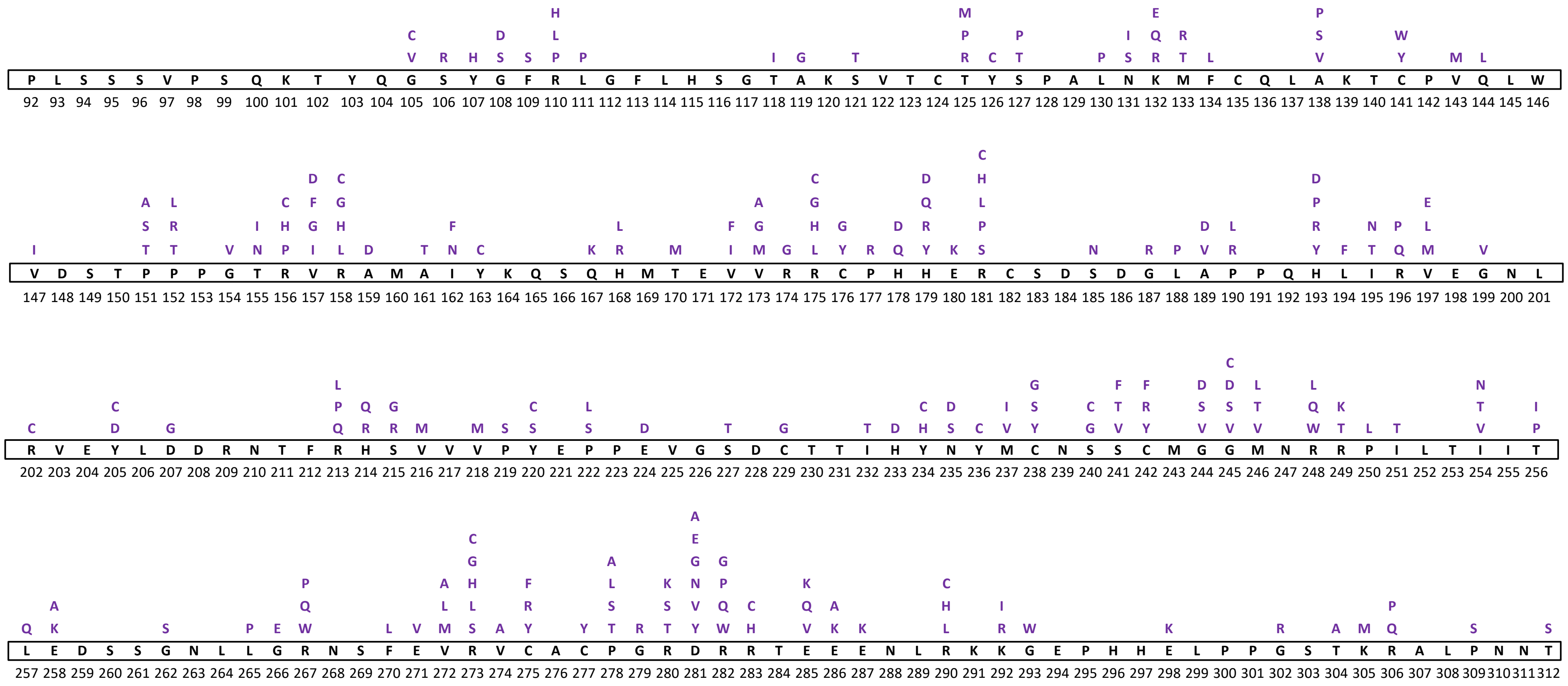


**Figure S1: Germline mutation map of p53 DBD.** The amino acid residues are represented by single-letter amino acid codes. The black color residues represent the wild-type residues whereas the violet color residues represent the mutations.
