## Supplementary material for "Phenotypical mapping of TP53 unique missense mutations spectrum in human cancers": Table S1

| **Residue** | **Residue position** | **Mutations reported so far in human cancers** | **Possible PTM** | **PTMs present in different pathways** | | | | | | | | |
| --- | --- | --- | --- | --- | --- | --- | --- | --- | --- | --- | --- | --- |
|  |  |  |  | **Transcriptional Activation** | **Transcriptional Suppression** | **DNA binding enhancement** | **Promoter specific DNA binding enhancement** | **Nuclear export** | **Nuclear export suppression** | **Degradation** | **Stabilization** | **Anti-repression** |
| **S** | **6** | **L** | **P** | **-** | **-** | **-** | **-** | **-** | **-** | **-** | **P** | **P** |
| **S** | **9** | **-** | **P** | **-** | **-** | **-** | **-** | **-** | **-** | **-** | **P** | **P** |
| **S** | **15** | **R** | **P** | **-** | **-** | **-** | **-** | **-** | **P** | **-** | **P** | **P** |
| **T** | **18** | **-** | **P** | **-** | **-** | **-** | **-** | **-** | **P** | **-** | **P** | **P** |
| **S** | **20** | **-** | **P** | **-** | **-** | **-** | **-** | **-** | **P** | **-** | **P** | **P** |
| **S** | **33** | **F, T** | **P** | **-** | **-** | **-** | **-** | **-** | **-** | **-** | **P** | **P** |
| **S** | **36** | **L, Q** | **P** | **-** | **-** | **-** | **-** | **-** | **-** | **-** | **-** | **-** |
| **S** | **37** | **P, T** | **P** | **-** | **-** | **-** | **-** | **-** | **-** | **-** | **P** | **P** |
| **S** | **46** | **F, P** | **P** | **-** | **-** | **-** | **P** | **-** | **-** | **-** | **P** | **P** |
| **T** | **55** | **-** | **P** | **-** | **-** | **-** | **-** | **-** | **-** | **-** | **-** | **-** |
| **T** | **81** | **I** | **P** | **-** | **-** | **-** | **-** | **-** | **P** | **-** | **P** | **P** |
| **K** | **120** | **E, M, N, Q, R** | **Ac** | **-** | **-** | **-** | **Ac** | **-** | **-** | **-** | **-** | **-** |
| **S** | **149** | **F, P, T** | **G, P** | **-** | **-** | **-** | **-** | **-** | **-** | **P** | **G** | **-** |
| **T** | **150** | **I, K, P, R** | **P** | **-** | **-** | **-** | **-** | **-** | **-** | **P** | **-** | **-** |
| **T** | **155** | **A, I, N, P, S** | **P** | **-** | **-** | **-** | **-** | **-** | **-** | **P** | **-** | **-** |
| **K** | **164** | **E, M, N, Q, R, T** | **Ac** | **-** | **-** | **-** | **Ac** | **-** | **-** | **-** | **-** | **-** |
| **S** | **215** | **C, G, I, N, R, T** | **P** | **-** | **-** | **-** | **-** | **-** | **-** | **-** | **-** | **-** |
| **E** | **258** | **A, D, G, K, Q, V** | **ADP** | **-** | **-** | **-** | **-** | **-** | **ADP** | **-** | **-** | **-** |
| **D** | **259** | **A, E, G, H, N, V, Y** | **ADP** | **-** | **-** | **-** | **-** | **-** | **ADP** | **-** | **-** | **-** |
| **E** | **271** | **A, D, G, K, Q, V** | **ADP** | **-** | **-** | **-** | **-** | **-** | **ADP** | **-** | **-** | **-** |
| **K** | **305** | **E, M, N, R, T** | **Ac** | **-** | **-** | **-** | **-** | **-** | **-** | **-** | **-** | **-** |
| **S** | **313** | **C, I, N, R** | **P** | **-** | **-** | **-** | **-** | **-** | **-** | **-** | **-** | **-** |
| **S** | **314** | **F** | **P** | **-** | **-** | **-** | **-** | **-** | **-** | **-** | **-** | **-** |
| **S** | **315** | **C, P** | **P** | **P** | **-** | **-** | **-** | **-** | **-** | **-** | **-** | **-** |
| **K** | **319** | **E, N** | **U** | **-** | **-** | **-** | **-** | **U** | **-** | **-** | **-** | **-** |
| **K** | **320** | **N** | **Ac, N, U** | **-** | **N** | **-** | **Ac** | **Ac, U** | **-** | **-** | **-** | **-** |
| **K** | **321** | **R** | **N, U** | **-** | **N** | **-** | **-** | **U** | **-** | **-** | **-** | **-** |
| **R** | **333** | **-** | **M** | **-** | **-** | **-** | **M** | **-** | **-** | **-** | **-** | **-** |
| **R** | **335** | **G, H, L** | **DM** | **-** | **-** | **-** | **DM** | **-** | **-** | **-** | **-** | **-** |
| **R** | **337** | **C, H, L, P** | **DM** | **-** | **-** | **-** | **DM** | **-** | **-** | **-** | **-** | **-** |
| **K** | **351** | **N** | **U** | **-** | **-** | **-** | **-** | **U** | **-** | **-** | **-** | **-** |
| **K** | **357** | **-** | **U** | **-** | **-** | **-** | **-** | **U** | **-** | **-** | **-** | **-** |
| **S** | **366** | **A** | **P** | **-** | **-** | **-** | **-** | **-** | **-** | **-** | **-** | **-** |
| **K** | **370** | **Q** | **Ac, M, N, DM, U, PU** | **-** | **M, N** | **A c** | **DM** | **Ac, U** | **-** | **PU** | **Ac** | **Ac** |
| **K** | **372** | **-** | **Ac, M, N, U, PU** | **M** | **N** | **Ac** | **-** | **Ac, U** | **-** | **PU** | **Ac** | **Ac** |
| **K** | **373** | **-** | **Ac, N, DM, U, PU** | **-** | **N** | **Ac** | **DM** | **Ac, U** | **-** | **PU** | **Ac** | **Ac** |
| **S** | **376** | **A, T** | **P** | **-** | **-** | **-** | **-** | **-** | **-** | **-** | **-** | **-** |
| **T** | **377** | **-** | **P** | **-** | **-** | **-** | **-** | **-** | **-** | **-** | **-** | **-** |
| **S** | **378** | **-** | **P** | **-** | **-** | **-** | **-** | **-** | **P** |  | **-** | **-** |
| **K** | **381** | **-** | **Ac, U, PU** | **-** | **-** | **Ac** | **-** | **Ac, U** | **-** | **PU** | **Ac** | **Ac** |
| **K** | **382** | **-** | **Ac, M, DM, U, PU** | **-** | **M** | **Ac** | **DM** | **Ac, U** | **-** | **PU** | **Ac** | **Ac** |
| **K** | **386** | **-** | **S, Ac, U, PU** | **S** | **S** | **Ac** | **-** | **S, U** | **-** | **PU** | **Ac** | **Ac** |
| **T** | **387** | **-** | **P** | **-** | **-** | **-** | **-** | **-** | **-** | **-** | **-** | **-** |
| **S** | **392** | **L** | **P** | **-** | **-** | **-** | **-** | **-** | **P** | **-** | **-** | **-** |

**Table S1: Mutation profile of post-translational specific residues in p53. (The PTM abbreviation used in the table are P= Phosphorylation; Ac=Acetylation; U=Mono-ubiquitination; PU=Poly-ubiquitination; M=Methylation; S=SUMOlyation; DM=Di-methylation; ADP= ADP ribosylation; N=Neddylation; G=O-GlcNAcylation)**
