## Supplementary material for "Phenotypical mapping of TP53 unique missense mutations spectrum in human cancers": Table S2

**Table S2: Distribution of all p53 unique missense mutations in different cancer tissues**

| **Tissues** | **TAD** | **PRR** | **DBD** | **DBD-OD linker** | **OD** | **CTD** |
| --- | --- | --- | --- | --- | --- | --- |
| Accessory sinuses | 0 | 0 | 83 | 1 | 0 | 0 |
| Adrenal gland | 1 | 0 | 36 | 0 | 3 | 0 |
| Anus and anal canal | 0 | 0 | 3 | 0 | 0 | 0 |
| Base of tongue | 0 | 0 | 11 | 0 | 0 | 0 |
| Bladder | 5 | 11 | 351 | 4 | 7 | 1 |
| Brain | 5 | 10 | 353 | 1 | 3 | 2 |
| Bones, joints and articular cartilage of limbs/other and unsppecified sites | 1 | 0 | 96 | 0 | 2 | 0 |
| Breast | 7 | 8 | 460 | 2 | 16 | 1 |
| Bronchus and lung | 9 | 4 | 482 | 4 | 13 | 2 |
| Cervix uteri | 0 | 0 | 59 | 0 | 0 | 0 |
| Colon | 5 | 0 | 260 | 0 | 1 | 4 |
| Colorectum | 0 | 0 | 298 | 1 | 7 | 0 |
| Connective, subcutaneous and other soft tissues | 1 | 2 | 168 | 0 | 0 | 1 |
| Corpus uteri | 2 | 0 | 102 | 2 | 1 | 1 |
| Esophagus | 4 | 4 | 356 | 0 | 2 | 0 |
| Eye and adnexa | 0 | 0 | 15 | 0 | 0 | 0 |
| Floor of mouth | 0 | 0 | 47 | 0 | 0 | 0 |
| Female genital organs | 2 | 0 | 1 | 0 | 0 | 0 |
| Gallbladder | 0 | 0 | 59 | 0 | 0 | 0 |
| Gum | 0 | 0 | 46 | 0 | 0 | 0 |
| Head and neck | 2 | 2 | 186 | 0 | 4 | 0 |
| Heart, mediastinum and pleura | 0 | 0 | 10 | 0 | 0 | 0 |
| Hematopoietic and reticuloendothelial systems | 7 | 1 | 241 | 0 | 3 | 1 |
| Hypopharynx | 1 | 1 | 75 | 0 | 1 | 0 |
| Kidney | 0 | 1 | 67 | 0 | 3 | 0 |
| Larynx | 1 | 1 | 138 | 1 | 2 | 1 |
| Lip (excludes skin of lip) | 0 | 1 | 22 | 0 | 0 | 0 |
| Liver and intrahepatic bile ducts | 2 | 5 | 287 | 1 | 10 | 0 |
| Nasal cavity and middle ear | 8 | 1 | 103 | 0 | 0 | 0 |
| Nasopharynx | 0 | 0 | 22 | 0 | 1 | 0 |
| Lymph nodes | 1 | 5 | 245 | 1 | 2 | 0 |
| Meninges | 0 | 0 | 2 | 0 | 0 | 0 |
| Oropharynx | 1 | 2 | 87 | 0 | 2 | 1 |
| Other and ill defiend sites | 0 | 0 | 4 | 0 | 0 | 0 |
| Other and ill-defiend digestive organs | 0 | 0 | 1 | 0 | 0 | 0 |
| Other endocrine glands and related structures | 0 | 0 | 5 | 0 | 0 | 0 |
| Other and unspecified parts of urinary organs | 0 | 0 | 17 | 0 | 0 | 0 |
| Other and unspecified parts of biliary tract | 1 | 0 | 40 | 0 | 0 | 0 |
| Other and unspecified parts of mouth | 1 | 1 | 203 | 1 | 3 | 1 |
| Other and unspecified male genital organs | 0 | 0 | 1 | 0 | 0 | 0 |
| Other and ill-defined sites in lip, oral cavity and pharynx | 0 | 0 | 5 | 0 | 0 | 0 |
| Other and unspecified female genitals | 0 | 0 | 15 | 0 | 0 | 0 |
| Other and unspecified major salivary glands | 1 | 0 | 12 | 0 | 0 | 0 |
| Other and illdefined sites within respiratory system and intrathoracic organs | 0 | 0 | 11 | 0 | 0 | 0 |
| Other and unspecified parts of tongue | 0 | 0 | 85 | 0 | 0 | 0 |
| Ovary | 5 | 5 | 354 | 2 | 12 | 1 |
| Palate | 0 | 0 | 18 | 0 | 0 | 0 |
| Pancreas | 1 | 0 | 158 | 0 | 4 | 0 |
| Parotid gland | 3 | 2 | 15 | 0 | 0 | 0 |
| Penis | 0 | 0 | 8 | 0 | 0 | 0 |
| Peripheral nerves and autonomic nervous system | 0 | 0 | 35 | 1 | 3 | 0 |
| Placenta | 0 | 0 | 2 | 0 | 0 | 0 |
| Prostate gland | 2 | 3 | 160 | 1 | 3 | 0 |
| Pyriform sinus | 0 | 0 | 4 | 0 | 0 | 0 |
| Rectosigmoid junction | 0 | 0 | 25 | 0 | 0 | 0 |
| Rectum | 1 | 2 | 180 | 1 | 0 | 0 |
| Renal pelvis | 0 | 0 | 32 | 0 | 0 | 0 |
| Retroperitoneum and peritoneum | 0 | 0 | 24 | 0 | 2 | 0 |
| Skin (excludes skin of vulva, Skin of Penis, Skin of Scrotum) | 7 | 16 | 270 | 0 | 8 | 0 |
| Small intestine | 0 | 0 | 12 | 0 | 0 | 0 |
| Spinal chord, cranial nerves, and other parts of the central nervous system | 0 | 0 | 3 | 0 | 0 | 0 |
| Stomach | 6 | 5 | 259 | 0 | 2 | 0 |
| Testis | 0 | 0 | 14 | 0 | 0 | 0 |
| Thyroid gland | 1 | 0 | 52 | 0 | 0 | 0 |
| Tonsil | 0 | 0 | 9 | 0 | 0 | 0 |
| Thymus | 0 | 0 | 15 | 0 | 0 | 0 |
| Upper- urinary tract | 3 | 0 | 59 | 0 | 3 | 1 |
| Ureter | 0 | 0 | 14 | 0 | 0 | 0 |
| Uterus | 0 | 0 | 40 | 0 | 0 | 0 |
| Vulva | 0 | 0 | 52 | 0 | 1 | 0 |
| Vagina | 0 | 0 | 3 | 0 | 0 | 0 |
