## Supplementary material for "Phenotypical mapping of TP53 unique missense mutations spectrum in human cancers": Table S3

**Table S3: Distribution of p53DBD missense mutations in different cancer tissues**

| **Tissues** | **ST** | **F** | **PF** | **NF** | **Total** |
| --- | --- | --- | --- | --- | --- |
| **Accessory sinuses** | 0 | 12 | 25 | 46 | **83** |
| **Adrenal gland** | 0 | 7 | 9 | 20 | **36** |
| **Anus and anal canal** | 0 | 0 | 0 | 3 | **3** |
| **Base of tongue** | 1 | 0 | 0 | 10 | **11** |
| **Bladder** | 11 | 61 | 72 | 207 | **351** |
| **Brain** | 7 | 42 | 63 | 241 | **353** |
| **Bones, joints and articular cartilage of limbs/other and unsppecified sites** | 0 | 10 | 13 | 73 | **96** |
| **Breast** | 9 | 77 | 93 | 281 | **460** |
| **Bronchus and lung** | 13 | 86 | 94 | 289 | **482** |
| **Cervix uteri** | 1 | 12 | 15 | 31 | **59** |
| **Colon** | 5 | 42 | 53 | 160 | **260** |
| **Colorectum** | 4 | 43 | 55 | 196 | **298** |
| **Connective, subcutaneous and other soft tissues** | 2 | 30 | 40 | 96 | **168** |
| **Corpus uteri** | 0 | 14 | 12 | 76 | **102** |
| **Esophagus** | 8 | 60 | 67 | 221 | **356** |
| **Eye and adnexa** | 0 | 1 | 2 | 12 | **15** |
| **Floor of mouth** | 0 | 7 | 4 | 36 | **47** |
| **Female genital organs** | 0 | 0 | 0 | 1 | **1** |
| **Gallbladder** | 0 | 7 | 8 | 44 | **59** |
| **Gum** | 1 | 4 | 5 | 36 | **46** |
| **Head and neck** | 3 | 25 | 29 | 129 | **186** |
| **Heart, mediastinum and pleura** | 0 | 1 | 0 | 9 | **10** |
| **Hematopoietic and reticuloendothelial systems** | 3 | 24 | 49 | 165 | **241** |
| **Hypopharynx** | 1 | 7 | 6 | 61 | **75** |
| **Kidney** | 2 | 5 | 9 | 51 | **67** |
| **Larynx** | 3 | 14 | 21 | 100 | **138** |
| **Lip (excludes skin of lip)** | 0 | 3 | 3 | 16 | **22** |
| **Liver and intrahepatic bile ducts** | 3 | 47 | 45 | 192 | **287** |
| **Nasal cavity and middle ear** | 0 | 24 | 31 | 48 | **103** |
| **Nasopharynx** | 0 | 5 | 4 | 13 | **22** |
| **Lymph nodes** | 4 | 32 | 38 | 171 | **245** |
| **Meninges** | 0 | 0 | 2 | 0 | **2** |
| **Oropharynx** | 0 | 6 | 12 | 69 | **87** |
| **Other and ill defiend sites** | 0 | 0 | 2 | 2 | **4** |
| **Other and ill-defiend digestive organs** | 0 | 0 | 0 | 1 | **1** |
| **Other endocrine glands and related structures** | 1 | 1 | 0 | 3 | **5** |
| **Other and unspecified parts of urinary organs** | 1 | 0 | 3 | 13 | **17** |
| **Other and unspecified parts of biliary tract** | 0 | 5 | 4 | 31 | **40** |
| **Other and unspecified parts of mouth** | 3 | 33 | 39 | 128 | **203** |
| **Other and unspecified male genital organs** | 0 | 0 | 0 | 1 | **1** |
| **Other and ill-defined sites in lip, oral cavity and pharynx** | 0 | 0 | 1 | 4 | **5** |
| **Other and unspecified female genitals** | 0 | 1 | 0 | 14 | **15** |
| **Other and unspecified major salivary glands** | 0 | 5 | 2 | 5 | **12** |
| **Other and illdefined sites within respiratory system and intrathoracic organs** | 0 | 0 | 2 | 9 | **11** |
| **Other and unspecified parts of tongue** | 1 | 14 | 12 | 58 | **85** |
| **Ovary** | 5 | 47 | 48 | 254 | **354** |
| **Palate** | 0 | 1 | 4 | 13 | **18** |
| **Pancreas** | 0 | 15 | 29 | 114 | **158** |
| **Parotid gland** | 1 | 1 | 6 | 7 | **15** |
| **Penis** | 0 | 0 | 0 | 8 | **8** |
| **Peripheral nerves and autonomic nervous system** | 0 | 5 | 3 | 27 | **35** |
| **Placenta** | 1 | 0 | 1 | 0 | **2** |
| **Prostate gland** | 3 | 25 | 41 | 91 | **160** |
| **Pyriform sinus** | 0 | 0 | 1 | 3 | **4** |
| **Rectosigmoid junction** | 0 | 1 | 5 | 19 | **25** |
| **Rectum** | 0 | 24 | 34 | 122 | **180** |
| **Renal pelvis** | 0 | 3 | 6 | 23 | **32** |
| **Retroperitoneum and peritoneum** | 0 | 3 | 2 | 19 | **24** |
| **Skin (excludes skin of vulva, Skin of Penis, Skin of Scrotum)** | 8 | 56 | 62 | 144 | **270** |
| **Small intestine** | 0 | 2 | 1 | 9 | **12** |
| **Spinal chord, cranial nerves, and other parts of the central nervous system** | 0 | 0 | 1 | 2 | **3** |
| **Stomach** | 9 | 39 | 51 | 160 | **259** |
| **Testis** | 0 | 2 | 5 | 7 | **14** |
| **Thyroid gland** | 0 | 5 | 7 | 40 | **52** |
| **Tonsil** | 0 | 0 | 0 | 9 | **9** |
| **Thymus** | 1 | 2 | 2 | 10 | **15** |
| **Upper- urinary tract** | 1 | 5 | 10 | 43 | **59** |
| **Ureter** | 0 | 0 | 2 | 12 | **14** |
| **Uterus** | 0 | 3 | 2 | 35 | **40** |
| **Vulva** | 3 | 4 | 9 | 36 | **52** |
| **Vagina** | 0 | 0 | 0 | 3 | **3** |
