## Supplementary material for "Phenotypical mapping of TP53 unique missense mutations spectrum in human cancers": Table S4

**Table S4: Distribution of DBD-OD linker missense mutations in different cancer tissues**

| **Tissues** | **Functional** | **Non-functional** |
| --- | --- | --- |
| Bronchus and lung | 4 | 0 |
| Breast | 2 | 0 |
| Brain | 1 | 0 |
| Ovary | 2 | 0 |
| Liver and intrahepatic bile ducts | 1 | 0 |
| Other and unspecified parts of mouth | 1 | 0 |
| Bladder | 4 | 0 |
| Larynx | 1 | 0 |
| Peripheral nerves and autonomic nervous system | 1 | 0 |
| Accessory sinuses | 1 | 0 |
| Corpus uteri | 1 | 1 |
| Rectum | 1 | 0 |
| Bones, joints and articular cartilage of limbs | 1 | 0 |
| Colorectum | 1 | 0 |
| Lymph nodes | 1 | 0 |
| Prostate gland | 1 | 0 |
