## Supplementary material for "Phenotypical mapping of TP53 unique missense mutations spectrum in human cancers": Table S5

**Table S5: Distribution of OD missense mutations in different cancer tissues**

| **Tissues** | **Super trans** | **Functional** | **Partially functional** | **Non-Functional** |
| --- | --- | --- | --- | --- |
| Adrenal gland | 0 | 1 | 1 | 1 |
| Bladder | 0 | 5 | 1 | 1 |
| Brain | 0 | 1 | 1 | 1 |
| Bones, joints and articular cartilage of limbs/other and unspecified sites | 0 | 0 | 0 | 2 |
| Breast | 0 | 7 | 3 | 6 |
| Bronchus and lung | 1 | 4 | 4 | 4 |
| Colon | 0 | 1 | 0 | 0 |
| Colorectum | 0 | 2 | 4 | 1 |
| Corpus uteri | 0 | 1 | 0 | 0 |
| Esophagus | 0 | 2 | 0 | 0 |
| Head and neck | 0 | 3 | 1 | 0 |
| Hematopoietic and reticuloendothelial systems | 0 | 1 | 0 | 2 |
| Hypopharynx | 0 | 0 | 0 | 1 |
| Kidney | 0 | 2 | 0 | 1 |
| Larynx | 0 | 1 | 1 | 0 |
| Liver and intrahepatic bile ducts | 0 | 4 | 2 | 4 |
| Nasopharynx | 0 | 1 | 0 | 0 |
| Lymph nodes | 0 | 2 | 0 | 0 |
| Oropharynx | 0 | 1 | 0 | 1 |
| Other and unspecified parts of the mouth | 2 | 1 | 0 | 0 |
| Ovary | 0 | 5 | 3 | 4 |
| Pancreas | 0 | 3 | 0 | 1 |
| Peripheral nerves and autonomic nervous system | 0 | 2 | 1 | 0 |
| Prostate gland | 0 | 1 | 0 | 2 |
| Retroperitoneum and peritoneum | 0 | 1 | 0 | 1 |
| Skin (excludes skin of vulva, Skin of Penis, Skin of Scrotum) | 1 | 6 | 1 | 0 |
| Stomach | 1 | 0 | 1 | 0 |
| Upper- urinary tract | 0 | 3 | 0 | 0 |
| Vulva | 0 | 1 | 0 | 0 |
