## Supplementary material for "Phenotypical mapping of TP53 unique missense mutations spectrum in human cancers": Table S6

**Table S6: Distribution of RD missense mutations in different cancer tissues**

| **Tissues** | **Functional** | **Partial functional** |
| --- | --- | --- |
| Bladder | 0 | 1 |
| Brain | 2 | 0 |
| Breast | 1 | 0 |
| Bronchus and lung | 1 | 1 |
| Colon | 2 | 2 |
| Connective, subcutaneous and other soft tissues | 1 | 0 |
| Corpus uteri | 1 | 0 |
| Hematopoietic and reticuloendothelial systems | 1 | 0 |
| Larynx | 1 | 0 |
| Oropharynx | 0 | 1 |
| Other and unspecified parts of mouth | 0 | 1 |
| Ovary | 0 | 1 |
| Upper urinary tract | 1 | 0 |
