## Supplementary material for "Phenotypical mapping of TP53 unique missense mutations spectrum in human cancers": Table S7

**Table S7: Tissue distribution of TAD and PRR mutations.**

| **Tissues** | **Functionality** | | | | | | | | | | | |
| --- | --- | --- | --- | --- | --- | --- | --- | --- | --- | --- | --- | --- |
|  | **Super tran** | | | **Functional** | | | **Partial functional** | | | **Non functional** | | |
|  | **TAD1** | **TAD2** | **PRR** | **TAD1** | **TAD2** | **PRR** | **TAD1** | **TAD2** | **PRR** | **TAD1** | **TAD2** | **PRR** |
| **Adrenal gland** | **0** | **0** | **0** | **0** | **1** | **0** | **0** | **0** | **0** | **0** | **0** | **0** |
| **Bladder** | **0** | **0** | **0** | **1** | **4** | **11** | **0** | **0** | **0** | **0** | **0** | **0** |
| **Bones, joints and articular cartilage of other and unspecified sites** | **0** | **0** | **0** | **0** | **0** | **0** | **0** | **1** | **0** | **0** | **0** | **0** |
| **Brain** | **0** | **0** | **0** | **0** | **4** | **9** | **0** | **1** | **1** | **0** | **0** | **0** |
| **Breast** | **0** | **0** | **0** | **3** | **1** | **8** | **0** | **2** | **0** | **1** | **0** | **0** |
| **Bronchus and lung** | **0** | **0** | **0** | **5** | **2** | **3** | **0** | **1** | **1** | **0** | **1** | **0** |
| **Colon** | **0** | **0** | **0** | **3** | **2** | **0** | **0** | **0** | **0** | **0** | **0** | **0** |
| **Connective, subcutaneous and other soft tissues** | **0** | **0** | **0** | **0** | **1** | **2** | **0** | **0** | **0** | **0** | **0** | **0** |
| **Corpus uteri** | **0** | **0** | **0** | **1** | **0** | **0** | **0** | **0** | **0** | **0** | **1** | **0** |
| **Esophagus** | **0** | **0** | **0** | **1** | **2** | **3** | **0** | **0** | **1** | **1** | **0** | **0** |
| **Female genital organs** | **0** | **0** | **0** | **2** | **0** | **0** | **0** | **0** | **0** | **0** | **0** | **0** |
| **Head and neck** | **0** | **0** | **0** | **0** | **1** | **2** | **0** | **0** | **0** | **1** | **0** | **0** |
| **Hematopoietic and reticuloendothelial systems** | **0** | **0** | **0** | **4** | **0** | **1** | **0** | **2** | **0** | **0** | **1** | **0** |
| **Hypopharynx** | **0** | **0** | **0** | **0** | **1** | **1** | **0** | **0** | **0** | **0** | **0** | **0** |
| **Kidney** | **0** | **0** | **0** | **0** | **0** | **1** | **0** | **0** | **0** | **0** | **0** | **0** |
| **Larynx** | **0** | **0** | **0** | **1** | **0** | **1** | **0** | **0** | **0** | **0** | **0** | **0** |
| **Lip (excludes skin of lip)** | **0** | **0** | **0** | **0** | **0** | **1** | **0** | **0** | **0** | **0** | **0** | **0** |
| **Liver and intrahepatic bile ducts** | **0** | **0** | **0** | **1** | **1** | **5** | **0** | **0** | **0** | **0** | **0** | **0** |
| **Lymph nodes** | **0** | **0** | **0** | **0** | **1** | **4** | **0** | **0** | **0** | **0** | **0** | **1** |
| **Nasal cavity and middle ear** | **0** | **1** | **0** | **0** | **4** | **1** | **0** | **3** | **0** | **0** | **0** | **0** |
| **Oropharynx** | **0** | **0** | **0** | **0** | **0** | **2** | **1** | **0** | **0** | **0** | **0** | **0** |
| **Other and unspecified major salivary organs** | **0** | **0** | **0** | **1** | **0** | **0** | **0** | **0** | **0** | **0** | **0** | **0** |
| **Other and unspecified parts of biliary tract** | **0** | **0** | **0** | **0** | **1** | **0** | **0** | **0** | **0** | **0** | **0** | **0** |
| **Other and unspecified parts of mouth** | **0** | **0** | **0** | **1** | **0** | **1** | **0** | **0** | **0** | **0** | **0** | **0** |
| **Ovary** | **0** | **0** | **0** | **3** | **0** | **5** | **0** | **1** | **0** | **1** | **0** | **0** |
| **Pancreas** | **0** | **0** | **0** | **1** | **0** | **0** | **0** | **0** | **0** | **0** | **0** | **0** |
| **Parotid gland** | **0** | **0** | **0** | **0** | **3** | **2** | **0** | **0** | **0** | **0** | **0** | **0** |
| **Prostate gland** | **0** | **0** | **0** | **1** | **1** | **3** | **0** | **0** | **0** | **0** | **0** | **0** |
| **Rectum** | **0** | **0** | **0** | **0** | **0** | **2** | **1** | **0** | **0** | **0** | **0** | **0** |
| **SKIN (excludes Skin of vulva, Skin of penis, Skin of scrotum)** | **0** | **1** | **0** | **1** | **3** | **15** | **0** | **1** | **0** | **1** | **0** | **1** |
| **Stomach** | **0** | **0** | **0** | **4** | **1** | **4** | **0** | **0** | **1** | **0** | **1** | **0** |
| **Thyroid gland** | **0** | **0** | **0** | **0** | **0** | **0** | **0** | **0** | **0** | **0** | **1** | **0** |
| **Upper urinary tract** | **0** | **0** | **0** | **2** | **0** | **0** | **0** | **1** | **0** | **0** | **0** | **0** |
